# Improved fluorescent *Listeria* spp. biosensors for analysis of antimicrobials by flow cytometry

**DOI:** 10.1101/2022.03.30.486417

**Authors:** Sebastian J. Reich, Jonas Stohr, Oliver Goldbeck, Bastian Fendrich, Peter Crauwels, Christian U. Riedel

## Abstract

The global increase in antibiotic resistances of pathogenic microorganisms require identification and characterization of novel antimicrobials. Bacterial biosensors expressing fluorescent proteins such as pHluorin variants are suitable for high-throughput screenings. Here, we present *Listeria* spp. pH-sensitive biosensors with improved fluorescence for single cell analysis of antimicrobials by flow cytometry.

## Text

The increasing global challenges with (multi)drug resistant bacteria highlight the demand for novel antimicrobial compounds to treat life-threatening infections [1]. Despite this growing need for novel anti-infective agents, the number of new antibiotics on the market is steadily decreasing [2, 3]. A major bottleneck in the development of new antimicrobial drugs is the lack of rapid, cost-effective, and reliable screening tools for lead compound identification [4]. Recently, our group has developed live a biosensor of the food-borne pathogen *L. monocytogenes* for detection of antimicrobial compounds that kill target bacteria by pore formation and disruption of membrane integrity [5]. The biosensor is based on monitoring of intracellular pH by expression of the GFP-derivative pHluorin, which is characterized by two distinct excitation peaks that change in relative fluorescence intensities in response to pH [6]. These biosensors were successfully used to determine susceptibility of bacteria to the lantibiotic nisin, measure antimicrobial activity in supernatants for natural and recombinant producers of antimicrobial peptides, and screen a library of bacteria isolated from raw milk for producer of antimicrobials [7-9]. Similar to the previously published biosensor strain *L. monocytogenes* EGDe/pNZ-P_help_-pHluorin (*Lm* pHin), a new vector was constructed, in which the pHluorin gene was replaced with a gene for pHluorin2, a pHluorin derivative with enhanced fluorescence [10]. The backbone of pNZ44 [11] was linearized by restriction with *Bgl*II and *Pst*I to remove the p44 promoter. The strong, constitutive P_help_ promoter was amplified from pPL2*lux*P_help_ [12] using primers P_help__fw (TTTTTATATTACAGCTCCAATCATTATGCTTTGGCAGTTTATTC) and P_help__rv (CTTTACTCATGGGTTTCACTCTCCTTCTAC). The gene encoding pHluorin2 was obtained as a synthetic DNA fragment codon-optimized for *Listeria monocytogenes* by a commercial service provider (Eurofins Genomics) and amplified using primers pHin2LM_fw (GTAGAAGGAGAGTGAAACCCATGAGTAAAGGTGAAGAATTATTTAC) and pHin2LM_rv (AGTGGTACCGCATGCCTGCACTATTTATATAATTCATCCATACCATGTG). Vector backbone and PCR products were assembled in a single isothermal reaction as described elsewhere [13]. Relevant parts of the resulting plasmid pNZ-pHin2^*Lm*^ (**Figure 1A**) were verified by Sanger sequencing (Microsynth Seqlab) and correct plasmids were used to transform *L. monocytogenes* EGDe and *L. innocua* LMG 2785. Following successful transformation, biosensor strains carrying plasmids pNZ-P_help_-pHluorin or pNZ-pHin2^*Lm*^ were initially checked for fluorescence by imaging in an iBright™ FL1000 Imaging System (ThermoFisher Scientific) with fluorescence detection mode at 488 nm (**Figure 1B**).

**Figure 1:**
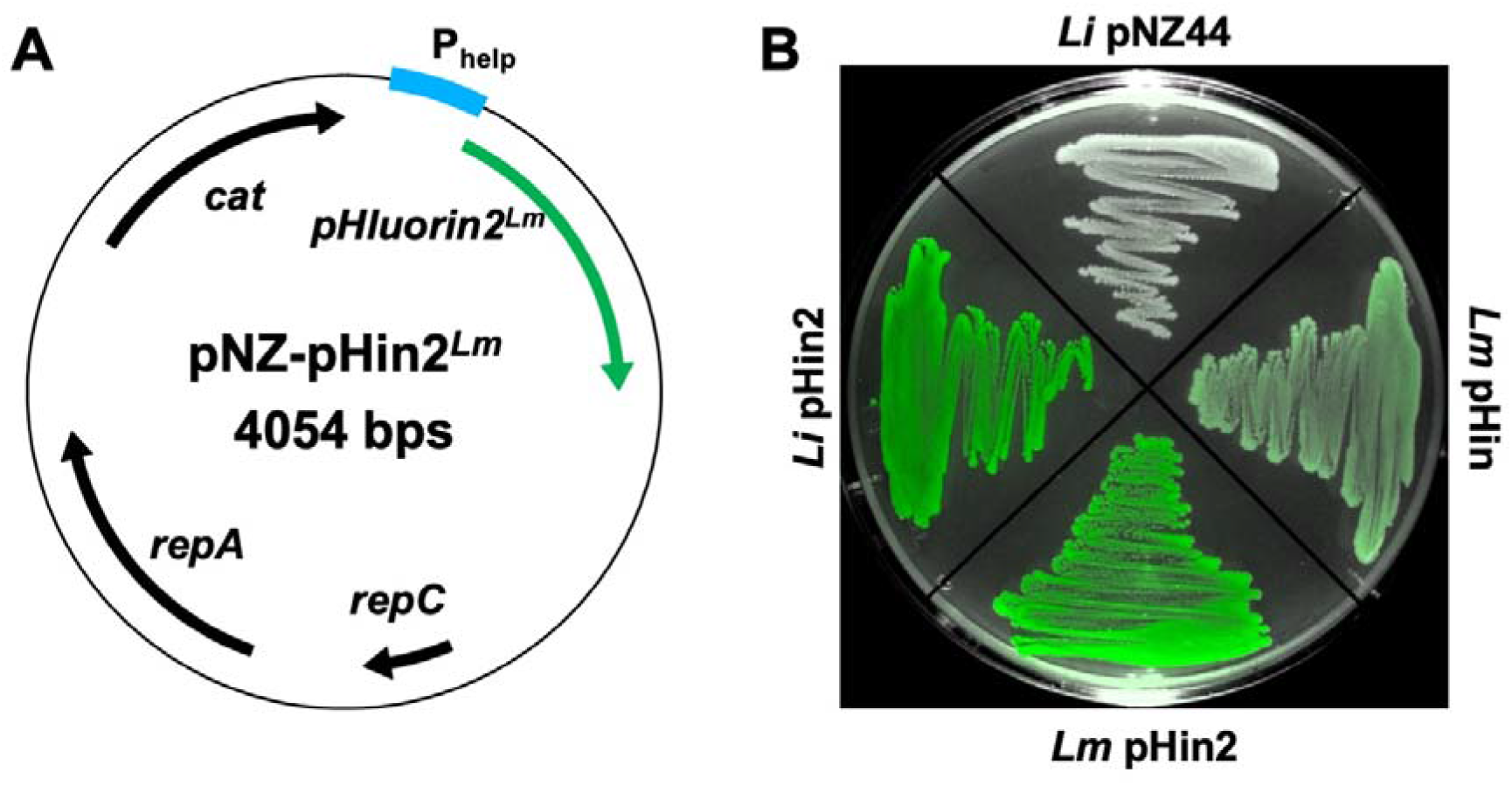
**(A)** Plasmid map of pNZ-pHin2^*Lm*^ and **(B)** fluorescence of *L. innocua* LMG 2785/pNZ44 (*Li* pNZ44), LMG 2785/pNZ-pHin2^*Lm*^ (*Li* pHin2), *L. monocytogenes* EGDe/pNZ-pHin2^*Lm*^ (*Lm* pHin2), or EGDe/pNZ-P_help_-pHluorin (*Lm* pHin) imaged in an iBright FL1000 in overlay (photograph and fluorescence detection mode at 488 nm).

Both new strains containing pNZ-pHin2^*Lm*^ (*Li* pHin2, *Lm* pHin2) showed brighter fluorescence on agar plates than the previously published strain *L. monocytogenes* EGDe/pNZ-P_help_-pHluorin (*Lm* pHin) whereas the empty vector control strain *L. innocua* LMG 2785/pNZ44 (*Li* pNZ44) showed no fluorescence above background. To further analyse fluorescence properties of the biosensors, bacteria were grown in BHI overnight (i.e. approx. 16 h), washed once in PBS and adjusted to an OD_600_ of 3 in filter-sterilized (pore-size 0.2 μm) LMB [5] adjusted to different pH (5.5-8.5). Aliquots of 100 μl were distributed into single wells of a black microtiter plate and mixed with 100 μl of LMB containing the cationic detergent cetyltrimethylammonium bromide (CTAB, final concentration 0.002% w/v). After incubation for 30 min at room temperature, fluorescence excitation spectra (350-490 nm) were recorded at an emission wavelength of 520 nm using a Tecan Infinite^®^ M200 multimode plate reader (Tecan, Crailsheim, Germany).

Similar to the previously published *Lm* pHin [5], *L. innocua* LMG 2785/pNZ-pHin2^*Lm*^ (*Li* pHin2) and *L. monocytogenes* EGDe/pNZ-pHin2^*Lm*^ (*Lm* pHin2) displayed the typical excitation spectrum of pHluorin proteins with excitation peaks at 400 and 475-480 nm (**Figure 2**). All three strains also showed the characteristic ratiometric, pH-dependent shift in fluorescence intensities across the excitation spectrum. However, fluorescence intensities were up to 6.7 and 9-fold higher for *Li* pHin2^*Lm*^ and *Lm* pHin2^*Lm*^ compared to *Lm* pHin depending on excitation wavelength and pH (**Figure 2)**. This is in line with data showing about 8-fold higher fluorescence for pHluorin2 over pHluorin when expressed in eukaryotic cells [10].

**Figure 2:**
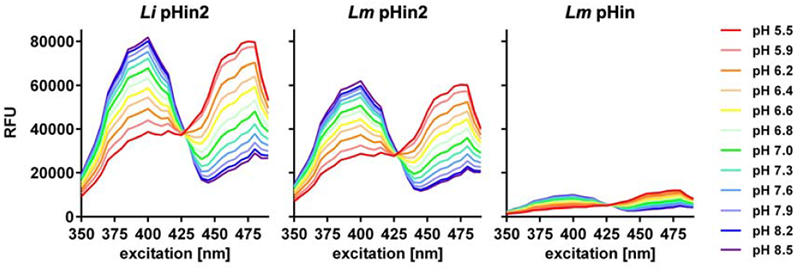
Relative fluorescence units at 520 nm (RFU) across a spectrum of excitation wavelengths (350-490 nm) of *L. innocua* LMG2785/pNZ-pHin2 (*Li* pHin2; left), *L. monocytogenes* EGDe/pNZ-pHin2 (*Lm* pHin2, middle) or EGDe/pNZ-P_help_-pHluorin (*Lm* pHin, right). Bacteria were resuspended in LMB adjusted to the indicated pH and permeabilized with CTAB (0.002%). Values are means of n = 3 independent cultures per strain.

To further demonstrate that the new biosensors behave similar to the previously published strain, dose-response experiment were performed with nisin A and pediocin PA-1, two antimicrobial peptides that kill target bacteria by disrupting membrane integrity [14, 15]. All three strains had comparable dose-response curves and showed a complete shift in excitation ratios at concentrations of 2.5 µg/ml of nisin and 625 ng/ml of pediocin, respectively (**Figure 3**). Furthermore, the pHin2 sensor strains showed a higher dynamic range of excitation ratios between ∼0.9 (for disrupted cells) to around ∼2.5 (intact), compared to ∼0.6 to ∼1.5 for pHin. This may be an advantage for measurements with high background fluorescence, e.g. supernatants of bacteria grown in complex media.

**Figure 3:**
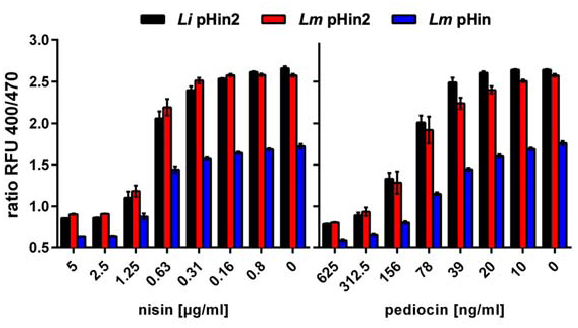
Ratios of fluorescence intensities at 510 nm after excitation at 400 and 470 nm (ratio RFU 400/470) of *L. innocua* LMG 2785/pNZ-pHin2 (*Li* pHin2; black bars), *L. monocytogenes* EGDe/pNZ-pHin2 (*Lm* pHin2, red bars) or EGDe/pNZ-P_help_-pHluorin (*Lm* pHin, blue bars) in LMB containing nisin (left) or pediocin (right) at the indicated concentrations. All values are mean ± standard deviation (SD) of n = 3 independent cultures per strain.

All three biosensor strains were further analyzed by flow cytometry using an Amnis^®^ CellStream^®^ device (Luminex) equipped with 405 and 488 nm lasers allowing excitation close to the two maxima of pHluorin proteins. For analysis of biosensor bacteria, flow speed was set to ‘slow’ and laser powers were 10% (forward scatter, FSC; side scatter, SSC), 35% (405 nm), and 40% (488 nm). Bacterial cells were identified based on FSC and SSC. Bacteria were then gated for singlet events based on the FSC aspect ratio. Analysis of singlets (min. 10 000 per sample) revealed that >98% of the bacteria showed bright fluorescence emission at 528 nm when excited with either the 405 or the 488 nm laser confirming homogenous expression of pHluorin2 by *Li* pHin2 (**Figure 4A**). Similar results were obtained for *Lm* pHin2 (data no shown). All three strains showed a single population with homogenous fluorescence in the 405/528 nm (excitation/emission) channel (**Figure 4B**). Flow cytometry confirmed about 8-fold higher fluorescence of *Li* pHin2 (38507 ± 2098 AU; N = 3 independent cultures) and *Lm* pHin2 (37886 ± 530 AU) in the 405/528 nm channel compared to *Lm* pHin (4143 ± 515 AU), which is in line with plate reader measurements (**Figure 2**). Additionally, flow cytometry was performed on untreated and nisin-treated (10 µg/mL, 30 min) *Li* pHin2 biosensors (**Figure 4C**) using PBS pH = 6.2 as sheath fluid. This allowed detection of biosensor bacteria in clearly distinct gates according to the 405/528 nm and 488/528 nm channels depending on the treatment. Bacteria in these two gates either represent untreated, alive/intact or nisin-treated, dead/disrupted bacteria. This demonstrates that flow cytometry can be used to assess intracellular pH and in consequence membrane integrity of pHluorin-expressing sensor bacteria on single cell level.

**Figure 4:**
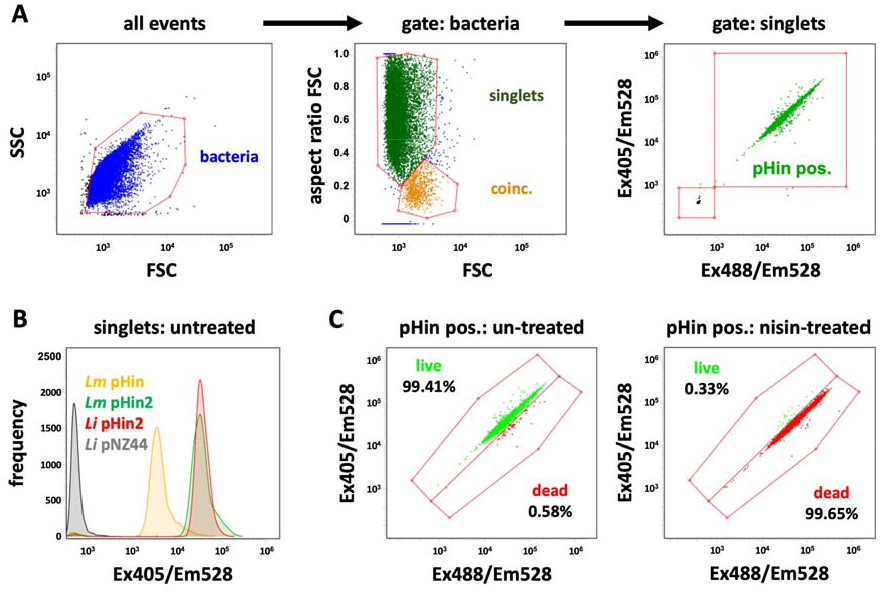
Analysis of *Listeria* spp. biosensor strains by flow cytometry. (**A**) Gating strategy to identify single bacterial cells that express pHluorin (pHin pos.). Amongst all recorded events, bacteria were gated based on their forward and side scatter (FSC, SSC; left panel). Singlets were identified by plotting the FSC aspect ratio over the FSC (middle) and pHluorin (pHin pos.) singlets were analyzed for fluorescence intensity (emission wavelength 528 nm) after excitation with the 405 nm (Ex405/Em528) and 488 nm laser right (Ex488/Em528). (**B**) Histogram plot of fluorescence intensity in the singlet gate of *Lm* pHin (yellow), *Lm* pHin2 (red), *Li* pHin2 (green), and the empty vector control strain *Li* pNZ44 (black). (**C**) Dot plots of fluorescence intensity of pHin positive events in the Ex405/Em528 and Ex488/Em528 channels for *Li* pHin2 after incubation with nisin (10 µg/mL, 30 min; right panel) or untreated controls (left panel). Results are representative of N = 3 independent cultures.

In conclusion, we provide two new biosensors of the genus *Listeria* that allow analysis of membrane damage using the ratiometric pH-dependent fluorescent protein pHluorin2 [10]. Both strains show up to 9-fold higher fluorescence compared to previously published strain *L. monocytogenes* EGDe/pNZ-P_help_-pHluorin [5]. All three strains behave comparable regarding challenge with membrane-damaging chemicals and peptides. The improved fluorescence properties of the new strains may facilitate analysis in matrices with high background fluorescence. Moreover, they were shown to be suitable for assessment of membrane integrity on single cell level using flow cytometry.

## Author contributions

Conceptualization: S.J.R. and C.U.R.; Methodology: S.J.R., O.G., P.C. and C.U.R.; Validation: S.J.R., O.G., P.C., and C.U.R.; Formal analysis: S.J.R., J.S., and B.F.; Resources: C.U.R.; Data curation: S.J.R., C.U.R.; Writing—original draft preparation: S.J.R. and C.U.R.; Writing—review and editing: S.J.R., OG and C.U.R.; Visualization: S.J.R. and C.U.R.; Supervision: S.J.R., O.G., P.C. and C.U.R.; Project administration: C.U.R.; Funding acquisition: C.U.R. All authors have read and agreed to the published version of the manuscript.

## Acknowledgements

This study was supported by a grant of the German Ministry for Education and Research to CUR (Grant No. 031B0826A) within the AMPLIFY consortium. The funding body had no role in the design of the study, analysis of the data, or writing of the manuscript.

## Ethics statement

None required.

## References

1. World Health Organization. Antimicrobial resistance: global report on surveillance. 2014.

2. Towse A, Hoyle CK, Goodall J, Hirsch M, Mestre-Ferrandiz J, Rex JH. Time for a change in how new antibiotics are reimbursed: Development of an insurance framework for funding new antibiotics based on a policy of risk mitigation. Health Policy. 2017;121(10):1025–30; doi: 10.1016/j.healthpol.2017.07.011.

3. Theuretzbacher U, Outterson K, Engel A, Karlén A. The global preclinical antibacterial pipeline. Nat Rev Microbiol. 2020;18(5):275–85; doi: 10.1038/s41579-019-0288-0.

4. Miethke M, Pieroni M, Weber T, Brönstrup M, Hammann P, Halby L, et al. Towards the sustainable discovery and development of new antibiotics. Nat Rev Chem. 2021;5(10):726–49; doi: 10.1038/s41570-021-00313-1.

5. Crauwels P, Schäfer L, Weixler D, Bar NS, Diep DB, Riedel CU, et al. Intracellular pHluorin as Sensor for Easy Assessment of Bacteriocin-Induced Membrane-Damage in Listeria monocytogenes. Frontiers in Microbiology. 2018;9:3038; doi: 10.3389/fmicb.2018.03038.

6. Miesenböck G, De Angelis DA, Rothman JE. Visualizing secretion and synaptic transmission with pH-sensitive green fluorescent proteins. Nature. 1998;394(6689):192–5; doi: 10.1038/28190.

7. Weixler D, Berghoff M, Ovchinnikov KV, Reich S, Goldbeck O, Seibold GM, et al. Recombinant production of the lantibiotic nisin using Corynebacterium glutamicum in a two-step process. Microb Cell Fact. 2022;21(1):11; doi: 10.1186/s12934-022-01739-y.

8. Desiderato CK, Sachsenmaier S, Ovchinnikov KV, Stohr J, Jacksch S, Desef DN, et al. Identification of potential probiotics producing bacteriocins active against Listeria monocytogenes by a combination of screening tools. International Journal of Molecular Sciences. 2021;22(16); doi: 10.3390/ijms22168615.

9. Goldbeck O, Desef DN, Ovchinnikov KV, Perez-Garcia F, Christmann J, Sinner P, et al. Establishing recombinant production of pediocin PA-1 in Corynebacterium glutamicum. Metabolic Engineering. 2021; doi: 10.1016/J.YMBEN.2021.09.002.

10. Mahon MJ. pHluorin2: an enhanced, ratiometric, pH-sensitive green florescent protein. Advances in Bioscience and Biotechnology. 2011;02(03):132–7; doi: 10.4236/abb.2011.23021.

11. McGrath S, Fitzgerald GF, van Sinderen D. Improvement and Optimization of Two Engineered Phage Resistance Mechanisms in Lactococcus lactis. Applied and Environmental Microbiology. 2001;67(2):608–16; doi: 10.1128/AEM.67.2.608-616.2001.

12. Riedel CU, Monk IR, Casey PG, Morrissey D, O’Sullivan GC, Tangney M, et al. Improved luciferase tagging system for Listeria monocytogenes allows realtime monitoring in vivo and in vitro. Applied and environmental microbiology. 2007;73(9):3091–4; doi: 10.1128/AEM.02940-06.

13. Gibson DG, Young L, Chuang R-Y, Venter JC, Hutchison CA, Smith HO. Enzymatic assembly of DNA molecules up to several hundred kilobases. Nature Methods. 2009;6(5):343–5; doi: 10.1038/nmeth.1318.

14. Brötz H, Josten M, Wiedemann I, Schneider U, Götz F, Bierbaum G, et al. Role of lipid-bound peptidoglycan precursors in the formation of pores by nisin, epidermin and other lantibiotics. Molecular Microbiology. 1998;30(2):317–27; doi: 10.1046/j.1365-2958.1998.01065.x.

15. Chikindas ML, García-Garcerá MJ, Driessen AJM, Ledeboer AM, Nissen-Meyer J, Nes IF, et al. Pediocin PA-1, a bacteriocin from Pediococcus acidilactici PAC1.0, forms hydrophilic pores in the cytoplasmic membrane of target cells. Applied and Environmental Microbiology. 1993;59(11):3577–84; doi: 10.1128/AEM.59.11.3577-3584.1993.

